# SplitThreader: Exploration and analysis of rearrangements in cancer genomes

**DOI:** 10.1101/087981

**Authors:** Maria Nattestad, Marley C. Alford, Fritz J. Sedlazeck, Michael C. Schatz

## Abstract

Genomic rearrangements and associated copy number changes are important drivers in cancer as they can alter the expression of oncogenes and tumor suppressors, create gene fusions, and misregulate gene expression. Here we present SplitThreader (http://splitthreader.com), an open-source interactive web application for analysis and visualization of genomic rearrangements and copy number variation in cancer genomes. SplitThreader constructs a sequence graph of genomic rearrangements in the sample and uses a priority queue breadth-first search algorithm on the graph to search for novel interactions. This is applied to detect gene fusions and other novel sequences, as well as to evaluate distances in the rearranged genome between any genomic regions of interest, especially the repositioning of regulatory elements and their target genes. SplitThreader also analyzes each variant to categorize it by its relation to other variants and by its copy number concordance. This identifies balanced translocations, identifies simple and complex variants, and suggests likely false positives when copy number is not concordant across a candidate breakpoint. It also provides explanations when multiple variants affect the copy number state and obscure the contribution of a single variant, such as a deletion within a region that is overall amplified. Together, these categories triage the variants into groups and provide a starting point for further systematic analysis and manual curation. To demonstrate its utility, we apply SplitThreader to three cancer cell lines, MCF-7 and A549 with Illumina paired-end sequencing, and SK-BR-3, with long-read PacBio sequencing. Using SplitThreader, we examine the genomic rearrangements responsible for previously observed gene fusions in SK-BR-3 and MCF-7, and discover many of the fusions involved a complex series of multiple genomic rearrangements. We also find notable differences in the types of variants between the three cell lines, in particular a much higher proportion of reciprocal variants in SK-BR-3 and a distinct clustering of interchromosomal variants in SK-BR-3 and MCF-7 that is absent in A549.

## Introduction

Genomic instability is one of the hallmarks of cancer, resulting in widespread copy number changes and structural variants including chromosome-scale rearrangements^1, 2^. Genomic rearrangements have been previously identified as drivers in some cancers, but they are not well understood and are difficult to detect and characterize. Genomic rearrangements include large structural variants such as chromosomal fusions, translocations, deletions, inversions, and duplications^3, 4^, and here we define them as being any novel adjacencies of sequences more than 10 kbp apart in the genome. The terms variant, rearrangement, and long-range structural variant are all used here interchangeably to refer to structural variants that connect sequences that used to be more than 10 kbp apart. These long-range variants often accompany major copy number variations, including cancer-driving amplification of oncogenes such as AKT2, ERBB2, and MYC, each of which have been found to initiate or exacerbate cancer^5-7^.

Copy number variants and gene fusions are common drivers in cancer, and both of which are linked to rearrangements^8, 9^. While commonly analyzed separately using read depth or split-read type analyses, copy number changes and structural rearrangements are highly interrelated. For instance, a large deletion manifests as both a decrease in copy number and a link between the two sequences on either side of the deletion. Gene fusions are also caused by rearrangements, and have a profound importance in cancer biology. For example, the first gene fusion discovered in cancer was BCR/ABL, which resulted from a fusion of chromosomes 9 and 22 and was found to be a driving mutation in chronic myelogenous leukemia^10^. However, the available algorithms for identifying gene fusions do not have perfect specificity, and require a joint analysis of genomic and transcriptomic data to correctly analyze.

With the advent of high-throughput sequencing, it has become possible to detect rearrangement variants in cancer genomes, but the sheer complexity of rearrangements, which often include adjacencies between distant regions of a chromosome or even between unrelated chromosomes, makes them difficult to study. Even the type of a variant can be obscured by other overlapping variants, which could have moved or flipped its breakpoints before or after the variant occurred. This occurs because variant types are determined by paired-end and split-read variant callers by the strands of the reads at two breakpoints, meaning the direction in which split reads map at each breakpoint^11, 12^. For instance, if a deletion occurs on the edge of an inverted sequence, one breakpoint of the deletion will be flipped, resulting in the deletion (+-) itself being called as an inversion (++), in addition to the original inversion. It is also a challenge to consider the total impact of these variants rearranging the genome as genes, regulatory elements, and all other genomic features may be repositioned relative to each other. Rearrangement variant calls carry some hints about the new context of these elements, but there are few tools for detailed investigations of these rearrangements across the genome, let alone analyzing their impact on chromosomal context of genes and regulatory elements.

We present our new open-source SplitThreader system, which uses a set of novel algorithms to explore, analyze, and search through rearrangements in cancer genomes. We demonstrate the application of SplitThreader to three different cancer cell lines with whole-genome sequencing: publicly available Illumina paired-end sequencing in A549^13^ and MCF-7^14^ analyzed using the LUMPY algorithm, and PacBio long-read sequencing in SK-BR-3 (Nattestad *et al*, in preparation) analyzed using the Sniffles algorithm (Sedlazeck *et al*, in preparation).

As shown in Figure 1, these three cell lines all contain hundreds of major rearrangements, many of which are part of complex sets of overlapping variants **(Supplementary Note 3)**. We use SplitThreader to search for gene fusions, analyze copy number concordance, and investigate the various types of rearrangements in these three cancer cell lines.

**Figure 1.**
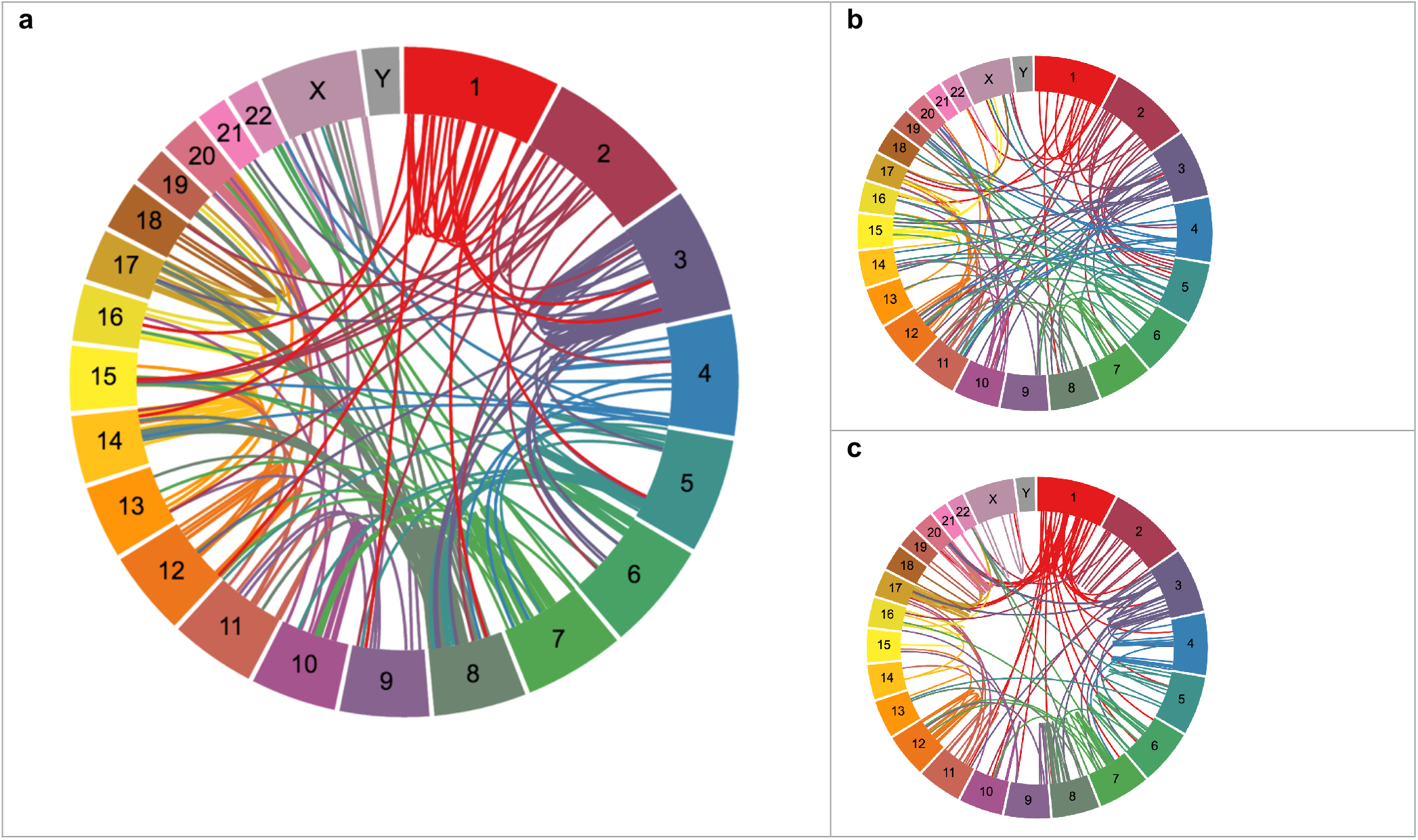
Circos plots showing genomic rearrangements in the cell lines SK-BR-3 (a), A549 (b), and MCF-7 (c).

### Searching through the landscape of rearrangements

Rearrangements in a cancer genome cause genomic features to be moved, reordered, or otherwise create a change in genomic context. For instance, the translocation, deletion, or any other repositioning of enhancers relative to a gene can be instrumental in defining the gene’s regulatory environment. Indeed, the *in vitro* insertion of a putative enhancer close to a gene was the method by which the enhancer was first discovered as a regulatory feature that can increase gene expression^15^.

In a complex set of hundreds of rearrangements taking place in some cancer genomes, systematically measuring and analyzing the movement of genes and regulatory features can be a challenge. To address this, SplitThreader builds a specialized sequence graph from the rearrangements in a genome, which encodes the positions of novel adjacencies introduced from the structural variations. Once defined, SplitThreader traverses the graph to analyze and calculate the new, post-rearrangement distances between genomic intervals in the sample. This graph search can be applied to many contexts, such as identifying the closest enhancer for each gene, the closest ChIP-seq peak to each transcriptional start site, or the closest evolutionarily conserved sequence to each insulator, to name a few possibilities.

Formally, the SplitThreader graph is composed of nodes representing sequences of DNA from a reference genome spanning between rearrangement breakpoints. The nodes have separate start and end ports so they can be read in either the forward or reverse complement directions: paths spanning from start to end will include the forward strand sequence, while paths spanning from end to start will include the reverse complement sequence. Edges in the graph are used to represent both rearrangement variants and non-rearranged reference-spanning connections between sequences. The edges are connected at each node to either the start or end port. The graph is initially constructed from a reference genome by creating a node for each chromosome. These nodes are then further split into separate nodes at all rearrangement breakpoints, with the appropriate edges introduced to maintain the original structure of the chromosome as well as the novel adjacencies created by rearrangements. SplitThreader always keeps the original reference allele edges because even if a variant-caller reports a homozygous variant, there can still be a low copy-number reference allele that is being outweighed by many copies of the variant allele. See the diagram in Figure 2 for an example of a small SplitThreader graph created from two chromosomes with two structural variants. The graph captures the possibilities for how the genomic sequence can be read, requiring that a node entered on one side must be exited on the opposite end, representing reading the sequence. This graph can then be traversed to calculate post-rearrangement distances between genomic intervals or find the closest genomic feature among a set of possibilities.

**Figure 2.**
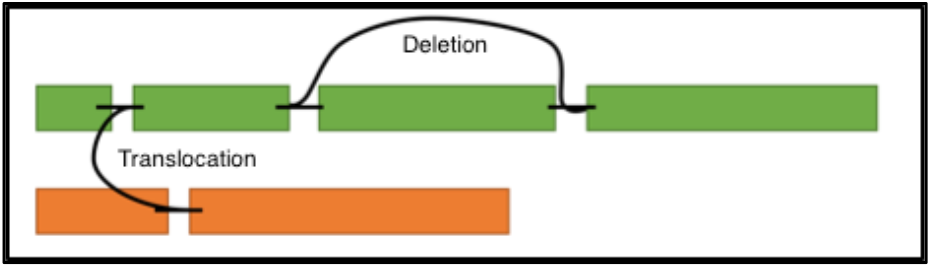
The structure of a SplitThreader graph representing a genome with two chromosomes (green and orange) and two rearrangements, one deletion and one inter-chromosomal translocation.

To perform a graph search, user specified target intervals, such as the coordinates of genes, enhancers, or other regulatory features, are marked in the graph, and then the ports for the starting intervals are added to a priority queue keyed by the genomic distance traversed. SplitThreader then uses the priority queue to perform a breadth-first search to find the shortest distance to a marked target port using a sequence graph-adapted implementation of the well-known Dijkstra’s algorithm. See **Methods** and **Supplementary Note 1** for a full description and diagrams of graph construction and priority queue breadth-first search. This general method allows users to find the shortest base pair distances between any set of intervals through the landscape of rearrangements in a genome.

### Finding rearrangements responsible for gene fusions

An important application of the SplitThreader graph search is to confirm and analyze gene fusions. RNA sequencing or other transcriptome evidence can provide a set of putative gene fusions, although the available methods often produce many false positives^16^. For this reason, gene fusion studies using RNA-seq often use PCR validation to confirm a transcriptome link between fusion genes^17, 18^. When whole-genome sequencing is available, it is possible to look for variants connecting the genes in the genome that were observed to be fused in the transcriptome, in a more high-throughput way than individual PCR validation. This can be done by manually looking for variants intersecting both genes, using genome arithmetic tools such as BEDTools pair-to-pair^19^, or using a dedicated fusion finder like BreakTrans^18^. However, there are many scenarios of gene fusions that have genomic evidence, yet their genomic evidence could not be found using these methods. In these cases there is no single variant that intersects both genes but instead the highly rearranged cancer genomes have “two-hop” (or greater) gene fusions, where the corresponding genomic regions are not directly fused to each other, but instead require passing through a third (or more) genomic region^18, 20^. For example, within the SK-BR-3 breast cancer cell line, there are two examples of two-hop gene fusions.

Even in these complex cases, SplitThreader can be used to search for genomic evidence of each putative gene fusion by searching for the shortest and lowest variant-count paths that connects the two genes in the rearrangement graph. Searching for genomic evidence of gene fusions is a special case of the SplitThreader graph search. By marking the ports near the first gene as the starting points and the ports near the second gene as the target end-points, the priority queue breadth-first search produces the shortest path in base pairs connecting the two genes. From previous literature, we identified 11 gene fusions in SK-BR-3 and 3 in MCF-7, all of which had been validated by PCR^17, 18^. Using SplitThreader, we found genomic evidence for 10 of 11 in SK-BR-3, two of which are two-hop gene fusions, and from MCF-7, we found genomic paths for all three. As a negative control, SplitThreader correctly finds no EML4-ALK gene fusion in A549, which has been used in studies of NSCLC as a negative control for this fusion. Figure 3 shows a selection of gene fusions with genomic paths found by SplitThreader, including both of the two-hop gene fusions in SK-BR-3, and **Supplementary Note 2** shows all gene fusions in detail.

**Figure 3.**
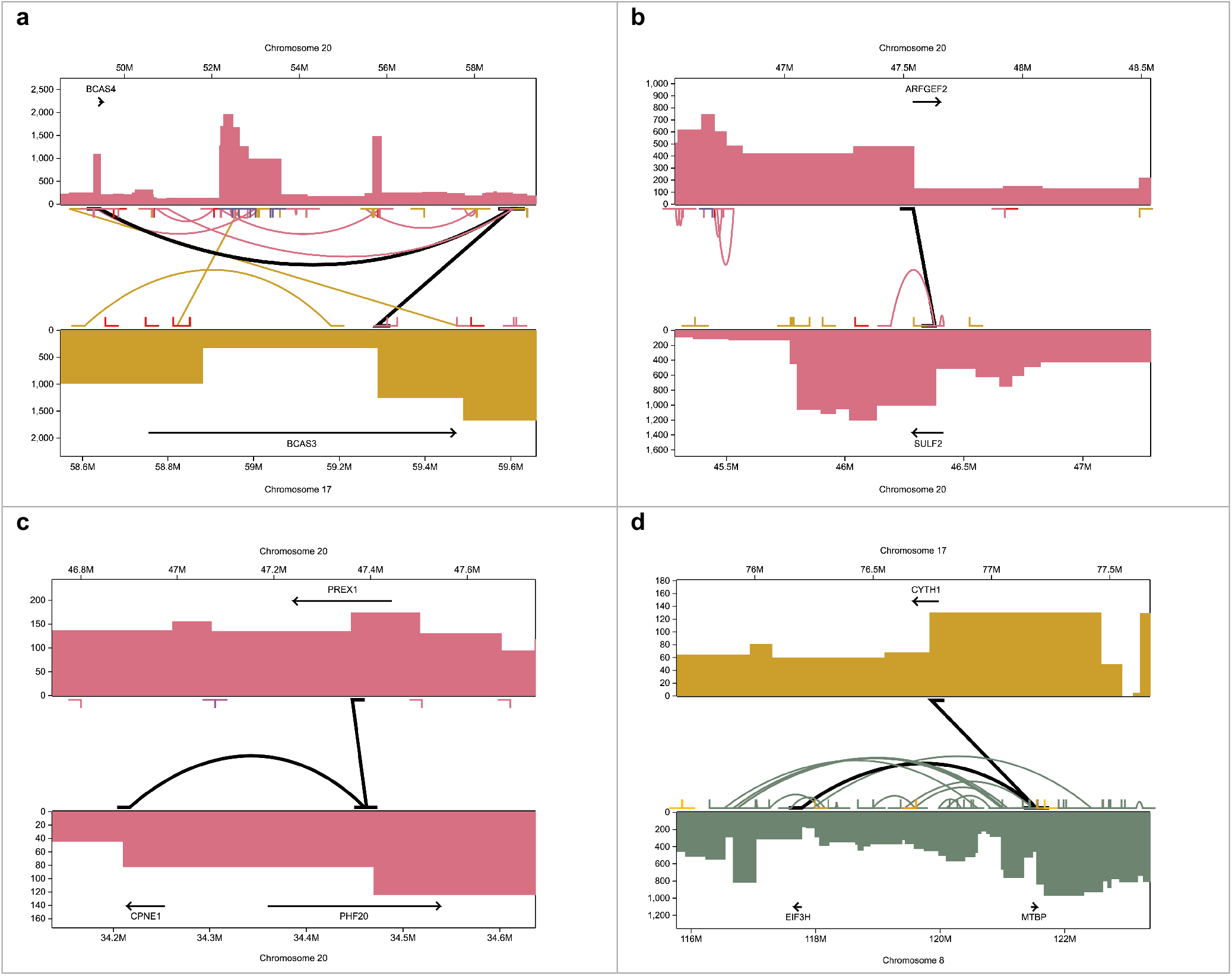
Selected gene fusions in MCF-7 and SK-BR-3 with genomic variant paths identified by SplitThreader. Segmented copy number profiles for up to two chromosomes are displayed as mirrored bar charts zoomed to the relevant regions. Variants are shown as connecting lines between the copy number profiles, and variants in the fusion path are shown as thicker black lines. The copy number profiles shown are segmented. (a) BCAS4-BCAS3 two-hop gene fusion gene fusion in MCF-7. (b) ARFGEF2-SULF2 gene fusion in MCF-7. (c) CYTH1-EIF3H two-hop gene fusion in SK-BR-3. (d) CPNE1-PHF20-PREX1 two-hop gene fusion in SK-BR-3.

### Variant neighborhood and copy number concordance analysis

Copy number changes are an extremely important form of variation in cancer genomes^8^, and their associated rearrangements give further insight into how these variants occurred. By combining copy number data and rearrangement variant calls from sequencing, SplitThreader can find the underlying rearrangements responsible for these copy number changes. In SplitThreader, each rearrangement variant is categorized by its copy number concordance (*matching, partial, non-matching*, or *neutral*) and variant neighborhood (*reciprocal, simple, solo*, or *crowded*). Diagrams representing each of the variant neighborhood categories intersected with each of the copy number concordance categories are shown in figure 4.

**Figure 4.**
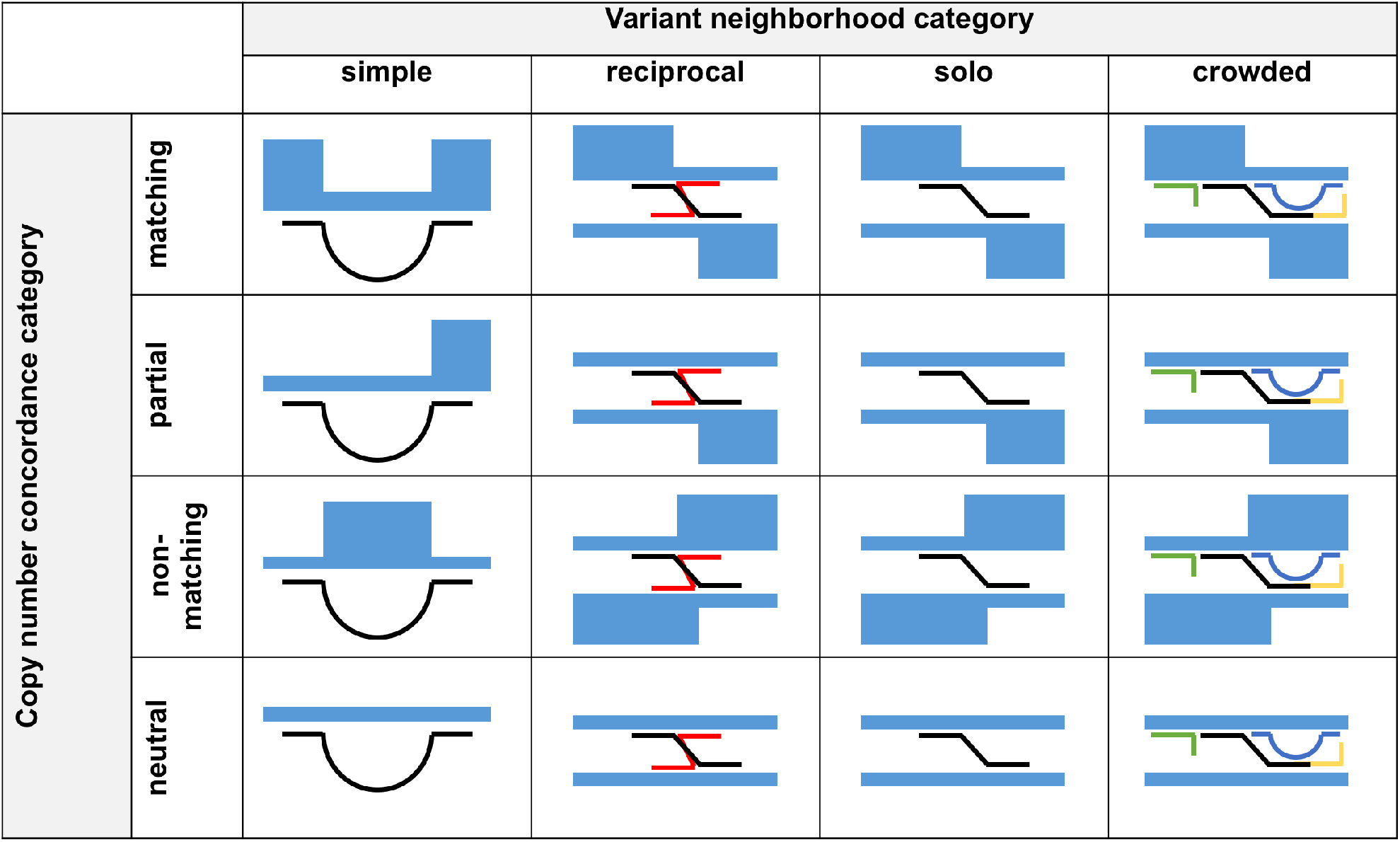
The variant categorization matrix with diagrams representing the intersections of the four variant neighborhood categories with the four copy number concordance categories.

*Simple* variants are defined by a lack of variants between their two breakpoints, which makes it likely that the type is robust and has not been obscured by rearrangements involving its breakpoints. *Solo* variants are at least 100 kbp away from other variants at both breakpoints, making the presence or absence of matching copy number changes more reliable to measure and attribute to that single variant. A *reciprocal* variant is a variant paired with another variant with each of its two breakpoints within 10 kbp of the breakpoints of the partner variant, and their strands are exactly the opposite, making the pair copy number neutral. Reciprocal variants can result from such events as translocations, mobile element insertions, and inversions. *Crowded* variants fit into none of the other categories, and therefore include most of the variants in complex, crowded, and overlapping regions.

These variant neighborhood categories define expectations for copy number concordance and classify variants for possible further analysis, such as statistics on manual curation and experimental validation. See **Methods** for details on how the categories are assigned. The intersection of the variant neighborhood and copy number concordance categorization schemes is especially important to prioritize manual curation. For instance, while reciprocal variants are expected to be copy number neutral, solo variants that are copy number neutral may be false positives. See **Supplementary Note 3** for examples of all variant neighborhood types and copy number concordance types, along with a detailed breakdown of how the category intersections can be interpreted. This categorization provides an overview of variant types and how copy number and rearrangements agree in a sample.

### SplitThreader is an interactive web application

All of the features discussed are accessible through the main SplitThreader web application available at http://splitthreader.com, which is open-source and can also be deployed locally from http://github.com/marianattestad/splitthreader. This includes copy number segmentation, categorization by copy number concordance versus variant neighborhood, and interactive queries through the nearest feature search and gene fusion search. Variants are shown in a table that can be filtered and sorted interactively. Variant counts are shown in a matrix of copy number concordance category versus variant neighborhood category, which can be used to inspect each variant in the category and even export any filtered subset of variants to a CSV file or to Ribbon^21^ for alignment visualization as part of further analysis of the variant calls. Gene fusion and graph search paths are found in real time and the results are immediately displayed in a table that can be used to inspect the rearrangements involved.

SplitThreader performs graph search, gene fusion finding, and categorization analysis on the fly in the browser, making it possible to use these features interactively to explore the data and the results. The visualization is created using D3^22^, a JavaScript library for linking data to visual elements in a web application. The visualization component in SplitThreader enables an overview of the raw data (Figure 5) and provides instant context for the results of the more complex algorithms such as graph search and gene fusion detection.

**Figure 5.**
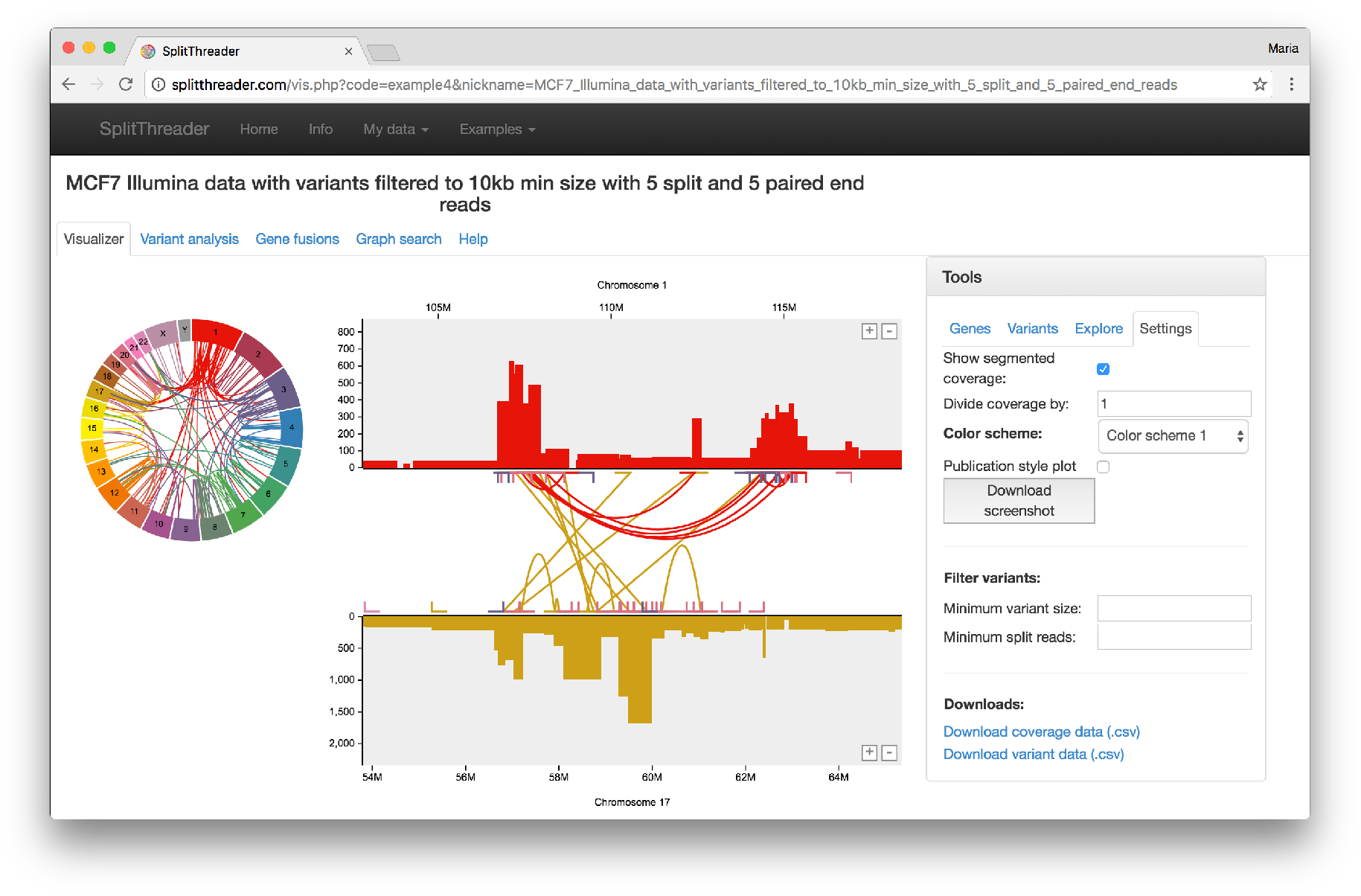
This screenshot of the MCF-7 results in SplitThreader shows the main visualization tab including a circos plot with rearrangements between all chromosomes, the copy number profiles of two selected chromosomes with rearrangements shown as connecting lines between them, and a panel with tools. Results of the general graph search, gene fusion search, and variant categorization analyses can all be inspected in this window.

## Discussion

SplitThreader provides a set of novel algorithms for searching through the landscape of rearrangements to calculate the new distances between genomic features, discover novel gene fusions, and explore how variants cluster together or mark changes in copy number. For example, within the cancer samples we analyze here we find that the SK-BR-3 and MCF-7 breast cancer cell lines both contain a few major clusters of intra- and interchromosomal variants, often stretching 5-20 Mb and containing anywhere from 5 to more than 100 variants. MCF-7 has three regions on chromosomes 1, 17, and 20 that each contain at least 20 variants connecting to the other two chromosomes. SK-BR-3 has a large region on chromosome 8 that contains 134 intrachromosomal and 66 interchromosomal translocations, 22 of which are connected to a cluster on chromosome 5. Using the variant neighborhood analysis, we also found that SK-BR-3 had a far higher proportion of reciprocal variants, 16.8% versus just 5.0% for MCF-7 and 3.1% for A549. Whether long-read sequencing has a higher sensitivity for reciprocal variants or SK-BR-3 actually more of these than the other two cell lines remains to be seen as more long-read sequencing is done in cancer in the coming years.

Categorization of variants based on the presence and relationship to nearby variants, known as variant neighborhood analysis, is an important tool for analyzing the qualities and types of variants. Copy number concordance analysis, especially when combined with variant neighborhood analysis, explains how rearrangements are supported by evidence from copy number variation. The intersection of these two categorization schemes provides new insights into how rearrangements may be disrupting normal gene regulation. The analyses also provide a strong starting point for manual curation and experimental validation, suggesting combinations of categories that are more or less likely to be true positives. To contextualize the results of these analyses, SplitThreader can be used to inspect each variant or path in the visualizer, showing copy number, rearrangements, and genes nearby. This combination of path-finding algorithms, manual curation features, and intuitive visualization makes SplitThreader an invaluable tool for inspecting rearrangements in a cancer genome.

## Methods

SplitThreader is comprised of one major data structure, the SplitThreader graph, and a core algorithm for traversing and searching through the graph, a specialized priority queue breadth-first search. In addition, SplitThreader uses related algorithms that for each variant determine its copy number concordance category and its variant neighborhood category. Here we discuss constructing the SplitThreader graph, performing priority queue breadth-first search across the graph, and evaluating nearest feature and gene fusion matches from the output of this algorithm. Then we explain in detail the methods for categorizing variants by their copy number concordance and their variant neighborhood.

### Constructing the SplitThreader graph

Nodes represent genomic sequences and edges represent connections between these sequences in the form of variants or original sequence in the reference. Each node has two ports, representing the start and end points of the sequence, and when traversing the graph, entering through one port requires exiting through the opposite port. If the start port is entered and the end port exited, this represents reading the sequence in the forward direction. If the end port is entered and the start port is exited, then this represents reading the sequence in the reverse complement direction. The graph is constructed as shown in **Supplementary Figure 1** where chromosomes are cut into nodes at the breakpoints of the variants and reconnected using edges to span the original reference and to represent the variants with the appropriate directionality to match the strandedness of the variants.

### Priority queue breadth-first search

SplitThreader uses a priority-queue breadth-first search strategy to find connections from a set of source intervals to a set of target intervals. A priority queue is used to prioritize shorter base-pair distance paths from source to target, such that the shortest base-pair distance solution is found first. The strategy consists of two preparatory steps followed by iteratively pulling from the priority queue until a solution is found.

First, the target locations are marked on the graph. The target intervals are overlaid on the graph to identify which nodes they overlap, and then each port on the overlapping nodes are marked with the ID of and distance to the target interval. If the port is inside the target interval, this distance will be zero, otherwise it is the distance in base pairs from the port to the start of the target interval within that node. The ID of the target interval is relevant for the nearest feature search where multiple features are marked as targets, and it is important to know which of these was found.

Second, the source locations are found and recorded with the same distance calculations as for the target ports, but now these are not marked onto the graph but instead are added to the priority queue where priority equals the distance from the source interval.

Third, an item is popped off the priority queue representing a port and its recorded path and cumulative distance so far, and the edges from the port are traversed. On the opposite side of each edge, we travel through the node (representing reading the sequence), and then add this port into the priority queue, recording its distance as the previous distance plus the length of the node we just traveled across. If the port we are adding to the priority queue is marked as a target, we add a target tag and record the distance as the cumulative distance so far plus the original target marker distance (recorded in the first step), and we add this to the priority queue. We repeat this third step, popping another port off the priority queue each time, until we pop off a target tag instead of a port, and then we have found the shortest distance from the source to the target.

This process finds the shortest path from any set of source intervals to any set of target intervals across the SplitThreader path. The strategy is outlined in **Supplementary Figure 2**.

### Evaluating graph search and gene fusion matches

This priority queue breadth first search (PQ BFS) is used in two different applications, a very flexible general graph search between any bed file intervals or genes, and a specialized gene fusion search.

The general graph search runs PQ BFS from each item in the source column against all items in the target column. A BED file of features can be uploaded, such as regulatory elements or repeats, which can be used as either the source, the target, or both (for instance finding the closest transcriptional start site for each enhancer). Thus it would be possible to use genes as the source and select only the lincRNAs, and then as the target upload a BED file containing regulatory features and select only the enhancers. Then running the calculation would produce for every lincRNA the path to the nearest enhancer.

Gene fusion search is done between any two genes in the annotation by name, so users can upload a list of paired gene names or enter pairs of gene names manually. PQ BFS is performed from a single source interval (gene 1) to a single target interval (gene 2), but this time the search is extended as far as 1 Mb distance even if matches are found earlier. All the matches are produced, and the best result by distance is reported for each number of variants. For example, if a 2-variant path is found with a distance of 152 kbp, the shortest 1-variant path found will also be reported, even if the distance is greater than 152 kbp.

### Copy number segmentation

SplitThreader starts its analysis with read alignment coverage information in 10 kbp bins. This can be generated from a sorted BAM file using a script called Copycat available on GitHub at https://github.com/marianattestad/copycat, which uses BEDTools^19^ genomecov to generate coverage for every base pair and then calculates averages into 10 kbp bins. The output of Copycat can be uploaded to SplitThreader, which then performs copy number segmentation using code adapted from Ginkgo^23^ that performs circular binary segmentation.

### Copy number concordance analysis

Whether a given rearrangement variant has a matching copy number change is informative for determining the accuracy of the variant call. The variant’s neighborhood of other variants influences whether a missing copy number change is indicative of a likely false positive or is consistent with expectations. For instance, reciprocal variants are expected to be copy number neutral, and variants close to each other can obscure the individual copy number changes produced by each variant. Because of this, copy number concordance analysis and variant neighborhood analysis should always be considered together. Here we outline the different categories of copy number concordance that variants can fall into. Each rearrangement variant has two breakpoints, for which we determine the closest copy number change from the segmentation. The copy number is considered concordant with a variant at a particular breakpoint if the copy number is higher on the same side of the breakpoint that the split reads are mapping. Copy number changes are only considered within 100 kbp of each breakpoint. “Matching” variants have concordant copy number changes on both breakpoints. “Partial” matching variants have a concordant copy number change on only one breakpoint. “Non-matching” variants have discordant copy number changes on one or both breakpoints. “Neutral” variants have no copy number changes within 100 kbp of either breakpoint. Each variant can only be in one copy number concordance category. The numbers of variants in these categories should be considered in combination with the categories in the variant neighborhood analysis.

### Variant neighborhood analysis

The variant neighborhood refers to a variant’s relationship with other variants nearby. A variant is classified as "reciprocal” if there is another variant that shares its two breakpoint locations (within 10 kbp) and has the opposite strands at both breakpoints. Some examples of variants that would show up as reciprocal are mobile element insertions, translocations, or copy number neutral inversions. “Simple” variants are intrachromosomal and do not have any overlapping variants between the two breakpoints. These tend to be smaller variants, and they are an important class because the variant types such as deletion, inversion, or duplication is not obscured by overlapping variants that may have moved or flipped breakpoints relative to when the event occurred. “Solo” variants have no other variants within 100 kbp of both their breakpoints. This type can be expected to show clear copy number concordance because no other variants are nearby to obscure the copy number. All variants that are not reciprocal, simple, or solo are then classified as “crowded,” indicating that they have overlapping variants and that there are other variants within 100 kbp. This type is the most difficult to curate for accuracy because lack of copy number concordance cannot be used to definitely dismiss the variant as inaccurate, given that other variants nearby could be creating opposing copy number changes. Each variant can only be in one variant neighborhood category. The variant neighborhood categories importantly interact with the copy number concordance categories. See **Supplementary Note 3** for examples of each category and an exploration of how we can interpret the intersection of different categories.

## Acknowledgements

This work was supported by NSF[DBI-1350041]; NHGRI [R01-HG006677].

